# Use of short-read RNA-Seq data to identify transcripts that can translate novel ORFs

**DOI:** 10.1101/2020.03.21.001883

**Authors:** Chaitanya Erady, Shraddha Puntambekar, Sudhakaran Prabakaran

## Abstract

Identification of as of yet unannotated or undefined novel open reading frames (nORFs) and exploration of their functions in multiple organisms has revealed that vast regions of the genome have remained unexplored or ‘hidden’. Present within both protein-coding and noncoding regions, these nORFs signify the presence of a much more diverse proteome than previously expected. Given the need to study nORFs further, proper identification strategies must be in place, especially because they cannot be identified using conventional gene signatures. Although Ribo-Seq and proteogenomics are frequently used to identify and investigate nORFs, in this study, we propose a workflow for identifying nORF containing transcripts using our precompiled database of nORFs with translational evidence, using sample transcript information. Further, we discuss the potential uses of this identification, the caveats involved in such a transcript identification and finally present a few representative results from our analysis of naive mouse B and T cells, human post-mortem brain and cichlid fish transcriptome. Our proposed workflow can identify noncoding transcripts that can potentially translate intronic, intergenic and several other classes of nORFs.

**One-line summary:** A systematic workflow to identify nORF containing transcripts using sample transcript information.

## Introduction

Broad segregation of the human genome into protein-coding and noncoding is dependent on evidence of translation from Open Reading Frames (ORFs) within these regions. Current annotated canonical ORFs consist of exons and introns bound by UTRs on either side. During the processing of pre-mRNAs into mRNAs, the introns are spliced out, leaving only the exons to be translated by cellular ribosomes. Noncoding regions on the other hand were initially termed “junk DNA”, but recent evidence has shown their ability to both transcribe and translate, ^1–4^. Additionally, several other novel transcripts and their functionality within the genome has been established ^5^. Moreover, contrary to the one gene-one protein hypothesis, multiple ORFs within known genes that can encode protein-products have been identified ^6,7^. Therefore, multiple translations from protein-coding and noncoding regions highlights that the human proteome is much more diverse than what is currently known and we call this separate and undefined class of open reading frames as novel open reading frames (nORFs).

Canonical ORFs code for the main protein from a protein-coding transcript whereas nORFs are as of yet unannotated and code for a different and in comparison a smaller protein. As shown in **Figure 1A**, nORFs can be found within the coding sequence (CDS) of known protein-coding genes, albeit in an alternative frame, within the UTRs, overlapping the UTR and CDS or antisense to known genes. nORFs can also be found within *de novo* genes, pseudogenes and other noncoding regions ^8^. Moreover, as shown in **Figure 1B**, nORF translation is observed across the human genome. Interestingly, chromosome Y has the lowest number of nORFs per unit 10,000 base pairs albeit on average the longest nORFs whereas chrMT is the most dense with short nORFs. The discovery of nORFs with evidence of translation renders evidence for a pool of unexplored proteins with potentially important functions some of which are discussed below.

**Figure 1:**
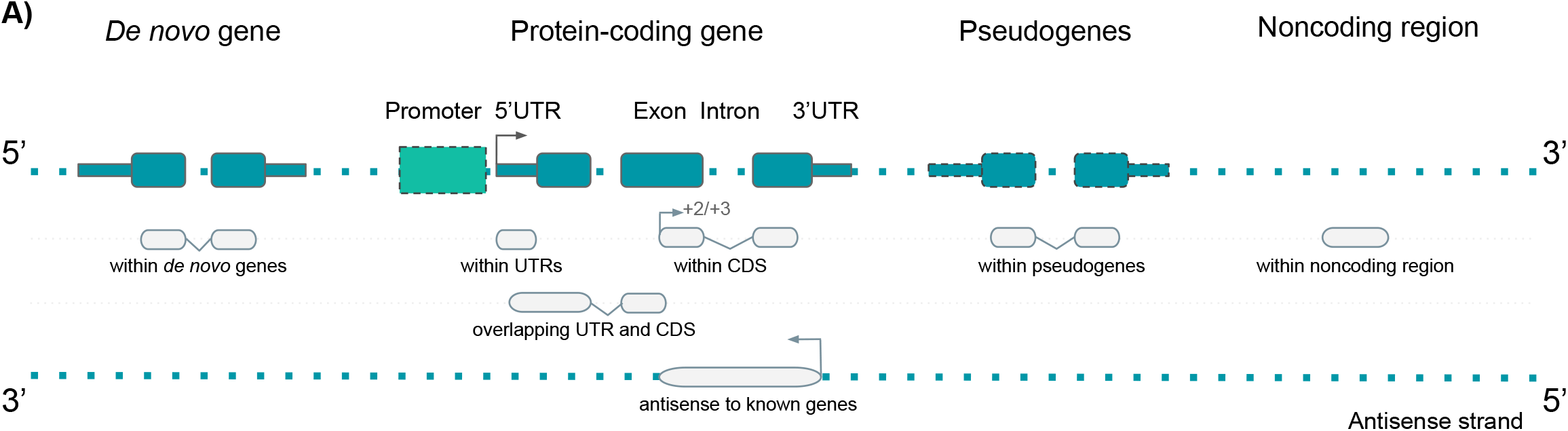
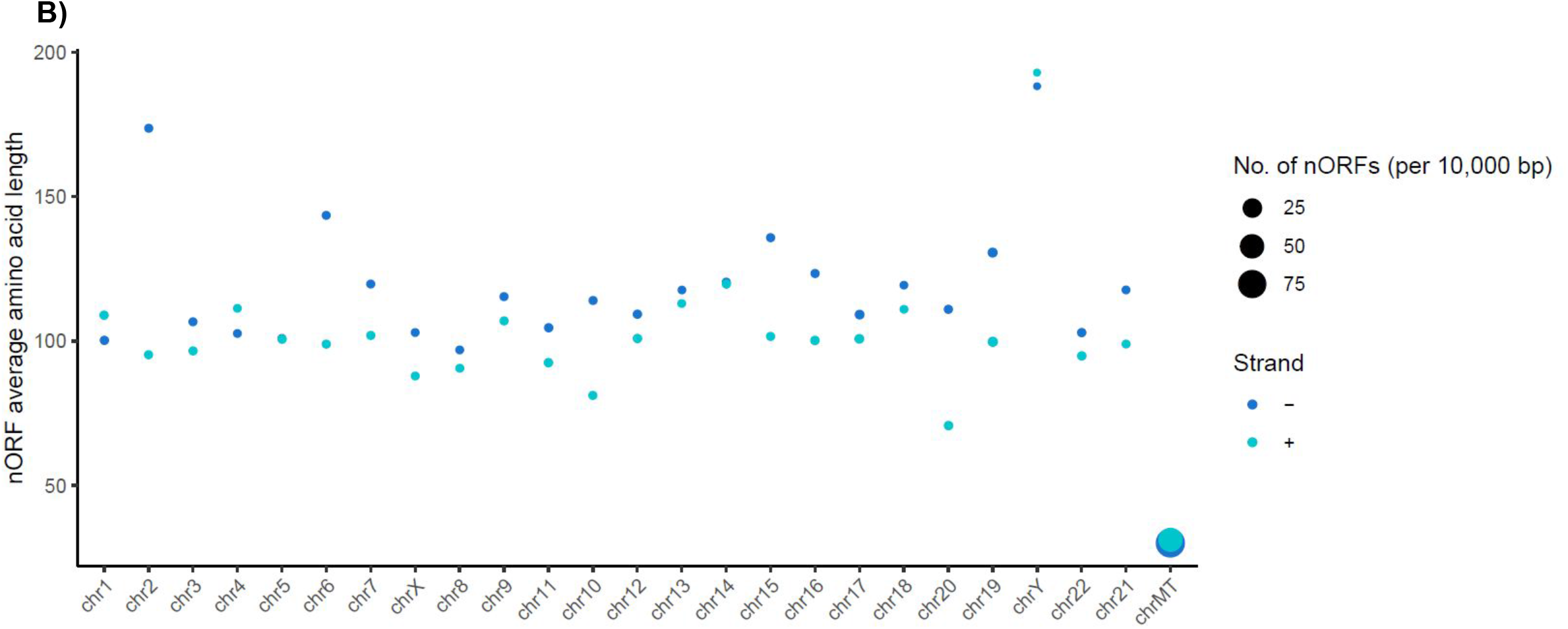
Novel ORFs (nORFs) in the human genome (GRCh38). **(A)** The different locations of nORFs (light grey boxes) within the genome is shown against known gene structures (blue-green boxes). nORFs can be present within the UTRs, the CDS but in an alternative frame, overlapping the UTRs and the CDS or antisense to known protein-coding genes. nORFs can also be found within de novo genes and pseudogenes as well as other noncoding regions. **(B)** Genomic distribution of nORFs in the human genome shows nORF expression is widespread. X-axis represents chromosome sorted according to decreasing size, y-axis shows average amino acid length of nORFs for a particular chromosome-strand pair, bubble sizes show no. of nORFs present within the chromosome-strand pair, normalised by chromosome size and scaled by a factor of 10,000 to give number of nORFs per 10,000 bp. chrY has the least density of nORFs whereas chrMT has the highest. Also, chrY on average, has the longest nORFs whereas chrMT has the smallest.

The presence of nORF encoded protein-like products, henceforth called as nORF products, and their utility has been explored in several species from bacteria to humans. To highlight a few-a 49-aa nORF product called AcrZ aids in *E. coli* survival against antibiotics by stimulating a drug expelling pump ^9^. Viruses diversify their proteome, given their small genomic sizes, using alternative ORFs ^10^. nORF products regulate plant morphogenesis and were also shown to regulate epidermal differentiation in *Drosophila* through the modification of a transcription factor called Shavenbaby ^11,12^. A mouse nORF called ‘Pants’, in a region orthologous to the 22q11.2 locus in humans, was identified with potential roles in hippocampal behaviour and a 68-aa protein called ‘NoBody’ in humans was found to manage the destruction of faulty mRNA in the cell by preventing the formation of P-bodies ^9,13^. Importantly, previous study from our lab demonstrated that nORF products from several noncoding regions in mouse neurons are not only translated but can also be biochemically regulated ^4^Λ More recently, we investigated nORF products in mouse B and T cells and showed that they can form structures, harbour deleterious mutations, and potentially be inhibited by drugs ^8^. These are just a few examples of why nORFs are significant genomic regions for further study.

The major hindrance in the identification and investigation of nORF products is their small size and lower abundance compared to canonical proteins ^8,14^. Moreover, existing gene prediction and computational methods for ORF predictions employ a minimum amino acid (aa) length threshold of 100aa, which filters nORFs from being detected ^15^. In addition, conventional gene signatures, for example, Kozak sequence and start codons, do not hold for nORFs, for which multiple start codons have been identified ^16^. Besides that, RNA-Seq experiments primarily employ poly-A-based selection of mRNAs, because of which almost 50% of the transcriptome which may not have poly-A, including noncoding RNAs such as miRNAs, lincRNAs and other nORF transcripts, are left undetected ^17^. Additionally, distinguishing between ORFs from the same region using RNA-Seq reads and detection of small protein products using current proteomic-based approaches is challenging ^18^. As a result most of the nORF products have been missed.

Recently, there have been a few computational and experimental attempts to overcome the above difficulties in identifying and investigating this unknown world of small proteins. One such computational tool is sORF finder, which aims to identify a class of very small nORFs (<= 100aa)^11^. Another example is mRNA assembler for proteogenomics (MAPS), that aims to improve transcriptome assembly for downstream proteogenomic analysis by using read sequences to build a consensus sequence database thereby supplanting annotated genome sequences for transcript assembly ^19^. Recent developments in experimental strategies to identify nORFs include Ribo-Seq, wherein transcripts that are bound to ribosome are sequenced under the assumption that these transcripts are translated ^18^, but the real evidence for translation from nORFs can only be obtained from mass spectra-based proteomics approaches. However, the current workflow in proteomics is limited because mass-spectra (MS) is only searched against a database of known proteins (usually from Uniprot or SwissProt). To identify nORF products, our lab and few others have pioneered a methodology modifying the current proteomics approach to a proteogenomic workflow. In this workflow (**Figure 2**), instead of searching the MS against a generic species-specific protein database, a custom database is made using the transcriptome or genome from a specific tissue and the MS from the same sample is searched against this custom database in six frames ^4,19,20^.

**Figure 2:**
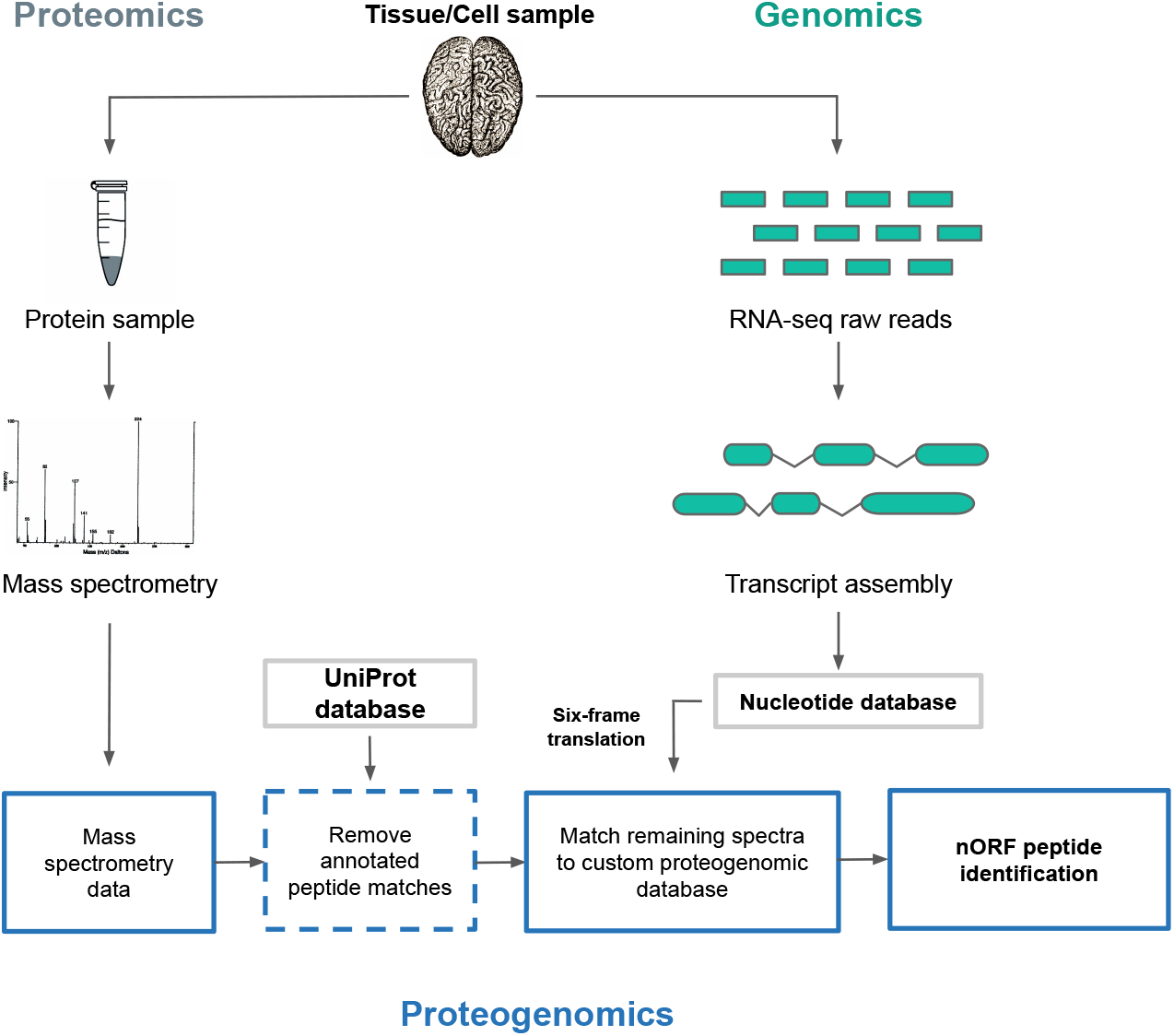
Proteogenomic workflow for nORF identification. Sample peptides identified using Mass spectrometry are first matched to UniProt database to remove known protein matches and then to a custom proteogenomic database to identify nORF peptides. RNA-seq data from the same samples used for protein collection is used for assembling transcripts which are then six-frame translated to produce the proteogenomic database.

Such improvements in technology have enabled attempts to curate evidence of nORF translation. For example, using evidence from computational prediction tools, ribosome profiling and literature mining, online resources such as OpenProt, sORFs.org, SmProt and ARA-PEP have been developed ^18,21–23^. But these attempts have remained sporadic and hence there has not been a single consistent definition for these unannotated nORFs. Therefore, we collated all the evidence published by multiple laboratories, re-classified the data and re-define them as nORFs ^14^. This curated data has been released in public domain available through https://norfs.org/ ^14^. This resource is expected to grow with constant updates from our research and other growing sources of nORF evidence. Through this we hope that nORFs can be systematically investigated as we did with nORFs.org entries where we noted a significant enrichment of deleterious mutations in these regions ^14^.

Besides all the above cited reasons, the current experimental strategies for nORF identification with Ribo-Seq or proteogenomics is prohibitive because of costs and for proteogenomic analysis it is imperative to obtain both RNA-Seq and MS from the same sample, which can be challenging. Therefore, in this study we present an alternative workflow using just RNA-Seq transcript data to identify transcripts translating nORFs curated in our continuously expanding nORF database. We discuss the caveats of using existing read-assembly, transcript-assembly and transcript-quantification pipelines for this purpose and propose a systematic workflow for nORF identification. Using our framework we can identify transcripts translating intronic and intergenic nORFs. Finally, we present results from the implementation of this workflow in mouse and human samples as proof-of-concept studies.

## Results

### Construction of custom databases for proteogenomic analysis

Here we describe experimental designs for three of our recent studies for identifying nORF products in mouse B and T cells, in human post-mortem brain samples (manuscript in preparation), and in cichlid fishes. We discuss in detail the pitfalls and the issues we overcame in the process.

For naive B and T cells isolated from the spleen of C57BL/6J mice, we assembled the RNA-seq reads into transcripts using HISAT2-StringTie (refer to materials and methods for details). Subsequently, we used bedtools getfasta ^24^ to extract transcript nucleotide sequence from the annotated reference genome, to create the custom proteogenomic database. Here, we encountered very long transcripts stitched together by the transcript assembler StringTie’s merge function ^25^. Such long transcripts were broken into lengths of less than 100,000 bps keeping with the search limit of 100,000 bps for Mascot (MS search engine). Although these transcripts may be technical artifacts, they were retained in our database to allow for any nORF identification present within them. MS of proteins isolated from these samples were first searched against the mouse UniProt database. To verify the presence of already known nORFs, the unmapped spectra, approximately 60%, were then mapped to a nORF amino acid database generated using the amino acid sequence of these nORFs obtained from sorfs.org and OpenProt ^18,21^,. Finally, the remaining unmatched spectra were matched to the custom proteogenomic database created using the sample-specific assembled transcriptome as shown in **Figure 2**. We used Mascot ^26^ for searching the spectra against the Uniprot proteins, the nORF amino acid database and the custom transcriptomic database in six frames, which is done “on the fly”.

Similarly, for the post-mortem brain samples, a combination of HISAT2-StringTie and bedtools getfasta were used to create the custom database and the protein MS from each sample was first matched against the human UniProt protein database and then any resulting unmatched spectra was mapped against the custom transcriptomics database for each sample. And for the cichlid study, HISAT2-StringTie assembled transcripts were clustered (k-means clustering) using Ballgown to reduce the number of highly similar assembled transcripts. Gffread utility (http://ccb.jhu.edu/software/stringtie/gff.shtml#gffread) was used to generate the transcriptomic fasta file which was searched against sample MS using Mascot, and proteins with at least 2 peptide matches were called at a false-discovery rate (FDR) less than 0.01.

### Caveats of the proteogenomic approach in identifying nORF protein products

The most prevalent method of identifying proteins is a “bottom-up” mass-spectrometry based approach. In this methodology, proteins are cleaved using proteases, separated using liquid chromatography columns, ionized and detected using a mass spectrometer. The resulting MS, is then computationally ‘matched’ to all theoretical MS of known proteins identified in that organism obtained from a pre-compiled database of known proteins like UniProt ^27^. Therefore, this approach allows only for the identification of already annotated proteins in that organism. In contrast, a proteogenomic approach, where the MS is searched against a custom database generated from a 3-frame or 6-frame translated version of either genomic or transcript sequences obtained from the same sample, is the only credential way to identify nORF encoded peptides ^4,20,28^. Additionally, using proteogenomics to identify peptide spectra and validate unannotated splice junctions and translations from regions currently annotated as noncoding, could aid in the refinement of existing genome annotations ^28,29^.

In preparing RNA-Seq reads for transcript assembly and custom proteogenomic database creation, the use of poly-A enriched transcripts, a dataset depleted of poly-A(-) and therefore several novel and noncoding transcripts as well as bimorphic transcripts ^30^, would limit nORF peptide detection. But, when using total-RNA datasets, one needs to keep in mind the presence of highly abundant rRNA in the sample, which could hinder the detection of low abundance RNAs. Therefore, for nORF peptide discovery using a custom database, the use of total RNA-Seq transcripts with rRNA depletion or using whole-genome sequencing (WGS) data is preferable. The custom proteogenomic database creation steps differ based on the type of data used. If a transcript dataset is used, their nucleotide sequences are 6-frame translated for custom proteogenomic database creation. In contrast, for WGS data, the genome is fragmented based on a specific fragment length, and each fragment is then 6-frame translated. Another potential issue with using genome or transcriptome based custom databases is the increased number of false-positives as detailed in Blakeley et al., 2012 ^31^. The size of the custom database is an important consideration as with increasing size, although the identification of peptides increases, the false positive rate, i.e. chances of incorrect peptide matches also increase ^31^. In order to evade the increase in false-positive rates, MS data is first mapped to known proteins in UniProt database, and then the unmatched spectra are mapped to the custom proteogenomic database as done by us previously in Prabakaran et al ^4^.

Moreover, since search engines use a decoy-target based approach to calculate false discovery rates (FDRs) of peptide-spectrum matches (PSMs), the enlarged search space because of six-frame translation with potentially noncoding frame sequences, undervalues true PSMs which are subsequently not called ^31^. Thus, filtering of potential noncoding frames prior to FDR calculations is recommended. Since our focus is on nORFs, such a filtering strategy may not be appropriate because conventional coding signatures do not apply for nORFs and therefore there are no applicable criteria for frame selection.

Although a proteogenomic approach offers an appropriate validation strategy for nORF identification, availability of only RNA-Seq data makes this approach difficult. Additionally, exploring the potential of nORFs to exert functions at the transcript rather than peptide level like lncRNAs ^3^, warrants a need for evaluation of nORFs at the transcript level. Therefore, we describe strategies for the same in the following sections.

### The perils of nORF identification using transcript data

To overcome the limitation in obtaining both total RNA-Seq based transcript and MS-based proteomic data from the same sample for proteogenomic analysis mainly because of the cost involved, we propose a simple workaround using existing validated information about nORFs. Information for 194,407 nORFs curated from online sources is available on nORFs.org (this database will undergo an update in the near future). Our method uses this curated knowledge and proposes a framework to infer nORF transcription in any human transcript dataset, as described below.

To identify nORFs with transcript data alone, we have to consider the following points. First, the genomic coordinates of the nORF should be encompassed by a transcript such that the codons that encode the nORF peptides are within the transcript. Second, the exons of the nORFs, which denote the nucleotide sequence that gets translated, must be encompassed within the exons of the transcript. This is because when the transcript is processed, the introns are spliced out and only the exons remain in the mature transcript. However, such detections are compounded by several transcripts originating from the same genomic region with overlapping exons and therefore mapping nORFs across them blurs the transcript identification process (**Supplementary Figure 1**). Additionally, such a lack of unique nORF containing transcript makes expression analysis like differential expression analysis between conditions particularly cumbersome.

A possible strategy to quantify nORF expression at the transcript level could involve mapping aligned reads from a BAM file to the nORF genomic coordinates and using the resultant counts for expression analysis. However, because of the small size of nORFs, sometimes even smaller than the read length, read count generation becomes an issue. Another major challenge in such an undertaking is the trouble in delineating reads that genuinely correspond to nORF expression versus reads that correspond to the more abundantly translated ORF within transcripts from the same genomic region. Transcript assemblers have encountered a similar issue in trying to quantify transcript isoforms in an environment of more abundant transcripts. For example, StringTie ^25^ uses a network flow algorithm first to construct the most abundant transcript, remove the reads associated with it and then continue constructing as many transcripts that can be explained by the remaining reads. This strategy works for transcript isoforms but may not be best for nORFs, which are not transcripts themselves and can be overlapped by several transcripts. We, therefore, resorted to using established methods of transcript expression value estimations generated by transcript assemblers to evaluate nORFs at the transcript level.

The choice of read aligners and transcript assemblers is another major factor that affects the number of transcripts identified and thereby nORF detection. Additionally, running these processes in de novo mode or with a reference genome also alters the transcript pool identified (**Supplementary Figure 2**). HISAT2 ^25^, TopHat ^32^ and STAR ^33^ read aligners were considered in our analysis. HISAT2 is preferred over TopHat as the former was designed to be a successor of TopHat and it is much faster and less memory intensive than TopHat ^25,34^. HISAT2 and STAR are both specialized at identifying novel splice sites which could aid in better downstream assembly of novel and unannotated transcripts within the samples and therefore lead to a more diverse transcriptome identification ^25,33^.

We evaluated several transcript aligners with reference and in de novo mode in our analysis for the cichlids study. Specifically, we evaluated StringTie ^25^ run with and without a reference genome, RSEM ^35^, Cufflinks run with and without a reference genome ^36^, Trinity ^37^ and MAPs ^19^. MAPs is a transcript aligner developed specifically for downstream proteogenomic analysis and uses read sequences to build a consensus sequence database and therefore does not use an annotated reference genome for transcript assembly ^19^. The result of this is transcripts assembled do not have an identifier, for example, an Ensembl transcript id, nor do they have count level estimates but only FPKM and TPM values for the transcripts. Moreover, these sample-specific transcripts were different across different samples, and without any ids, it was hard to compare across samples for downstream differential expression analysis. For the post-mortem brain study, we tried running StringTie -merge on MAPs output but this grossly overestimated the transcriptome from an average of 23,000 transcripts per sample to an average of 150,000 transcripts after StringTie -merge. As StringTie-merge creates a union database and the subsequent StringTie run quantifies the merged transcripts for each sample, the increase in number shows how the transcripts called by MAPs across samples are very different and this strategy, therefore, may not be beneficial for differential expression analysis. Thus, we did not use MAPs for our analysis. In our evaluation of StringTie, Cufflinks and Trinity, we found that Trinity generated a very high number of transcripts with lower sensitivity and precision than StringTie and Cufflinks (**Supplementary Figure 2 and 3**). Moreover, RSEM and StringTie are well-cited tools and have shown to be better than other assemblers for transcript assembly, especially lncRNAs, which is an important category in our investigation ^38–40^. StringTie, in comparison to Cufflinks, calls a higher number of correctly called transcripts and is tailored to detecting novel isoforms which is vital for nORF identification ^40^. Therefore, we have resorted to using HISAT2-StringTie or STAR-RSEM for read alignment and transcript assembly prior to nORF identification.

To identify nORFs within these assembled transcripts, we initially used bedtools intersect ^24^ to find overlap between nORFs and all known transcripts from Ensembl ^41^ based on their genomic coordinates. However, this method proved incorrect since we were calling transcripts which encompassed nORFs while allowing for nORF overlap with the transcript intron, meaning, the nORF sequence would not be processed into the mature transcript. We, therefore, use GffCompare ^25,42^, since it identifies nORF-transcript matches based on intron chain overlap and thereby ensures that the nORF exons are encompassed within the identified transcripts. We introduced additional processing steps on the GffCompare result to identify these transcripts as detailed in the next section.

### Workflow for identification of nORFs using transcript data

To identify nORFs within the assembled transcriptome, we use GffCompare ^42^, a tool designed for annotating, merging, comparing and estimating the accuracy of assembled transcripts. We specified our nORF dataset as the reference using the -r parameter, which was searched against the query sample transcripts. If an exact intron chain match is found between the nORF and the sample transcript, a class code of “=“ is assigned to the match; otherwise a class code “c” is assigned for partial matches. The output is obtained through the REFMAP file wherein for each entry in the reference transcript, query transcripts matches are provided. A point of consideration is that GffCompare is stringent about removing any duplicates within the reference transcript file. Therefore, some of the nORFs which are identified as duplicates will not be considered for mapping. In brief for single-exon transcripts, if the length of transcript 1 is at least 80% of transcript 2 and transcript 1 is encompassed within transcript 2, transcript 1 is considered duplicated ^42^. For multi-exon transcripts, if transcript 1 has complete intron chain match with transcript 2 and transcript 1 is fully contained within transcript 2, transcript 1 is considered duplicated ^42^. If nORF duplicates are identified and removed as described above, it still does not affect our analysis since the longer nORF would still be retained in the reference database for transcript identification.

GffCompare results in the REFMAP file were processed, as shown in **Figure 3**. First, only those matches with class code “=“, i.e. overlaps between nORFs and sample transcripts with exact intron chain matches are retained. Next, the terminal exon boundaries are filtered such that the nORF coordinates are equal to or less than the sample transcript’s exon coordinates. This filter is used to ensure that the nORF coordinate, which starts with the translation start of the nORF, is encompassed within the sample transcript. Finally, sample transcripts annotated as protein-coding are removed from further analysis. The last filter is in place because protein-coding transcripts already have a known ORF within them and therefore the transcript’s expression value may correspond more to the canonical ORF than the nORFs. In contrast, noncoding transcripts, by definition, do not have ORFs that are translated within them. nORFs with experimental validation of translation albeit from other samples, are the only identified ORFs within these noncoding transcripts, and therefore their transcript abundances can be assigned to the nORFs within them.

**Figure 3:**
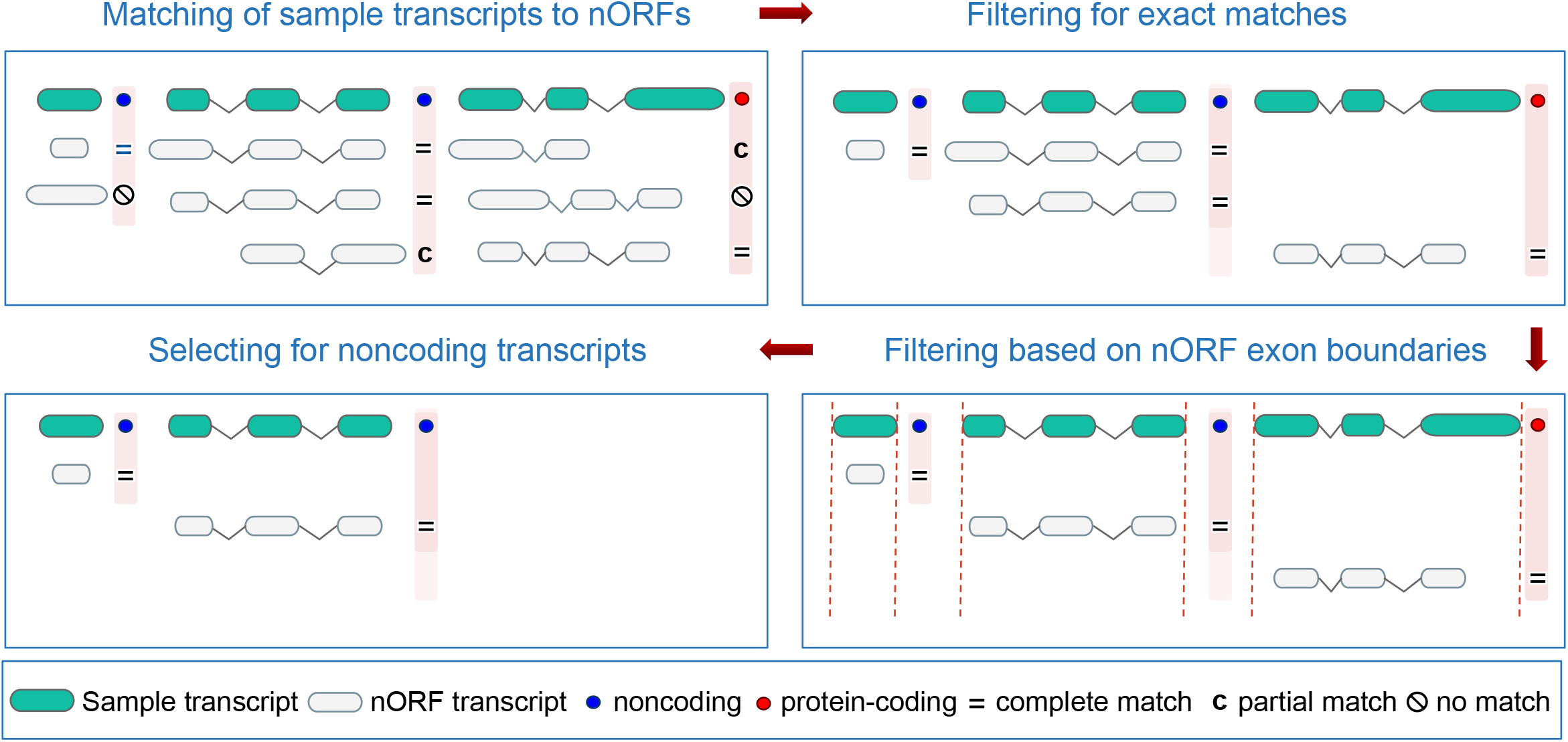
Workflow for nORF identification at the transcript level using. Using GffCompare, nORFs mapping to sample transcripts, based on intron chain matches are identified and assigned a classification code. The results are then filtered to retain only those matches with code “=“ representing complete match between the nORF and the transcript. Next, an exon boundary filter is imposed to ensure that the nORF exons are confined within the transcript exons. Finally, only sample transcripts with a non protein-coding biotype are retained for further analysis.

Based on nORF location within the genome and particularly the transcriptome, **Figure 4** highlights the different cases one might encounter, and which of these can be identified using the workflow and for expression analysis. Using our approach, we identify several nORFs within the noncoding genome. Additionally, using the prescribed workflow, we encounter cases where one transcript corresponds to multiple nORFs. This could highlight that multiple ORFs are being translated from the same transcript ^6^ or be a result of incorrect nORF identification in which case we cannot delineate which one is correct. Similarly, cases where one nORF matches to more than one transcript were also identified owing to the presence of multiple transcripts arising from the same region with similar intronic chain pattern and exon number. Again, using the current dataset, we cannot delineate and identify the “correct” nORF transcript. Usually ranked annotation schemes are used to select one transcript over the other ^18^ but we chose to process all identified noncoding transcripts for expression analysis. The workflow defined above allows for identification of nORFs and the corresponding transcript expression values for downstream analysis.

**Figure 4:**
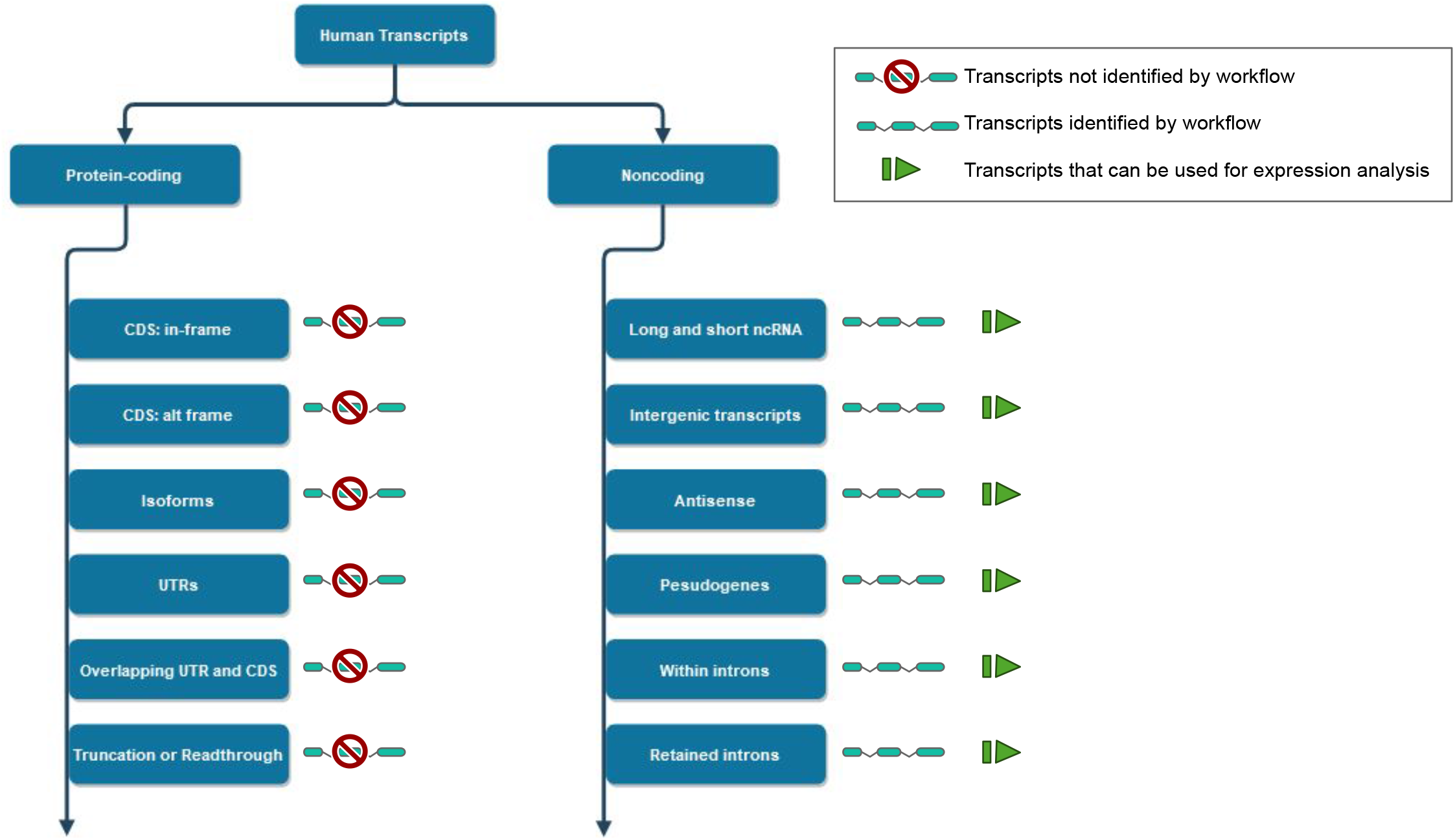
Types of transcripts that can be identified using the proposed workflow. The different types and sub-types of transcripts that can be identified using the workflow shown in Figure 3 as well as whether they are suitable for downstream expression analysis is displayed above.

### Proof-of-concept studies using our methodology

To demonstrate the use of the proposed workflow, in this section, we present examples from mouse immune and human brain samples. Mouse B and T cells were assembled using HISAT2-StringTie pipeline (see materials and methods for further details). Using the workflow described, we were able to identify nORFs and their transcripts and additionally were able to verify translation of a few nORFs using our proteogenomic workflow (**Figure 5A**). Moreover, examples of transcripts containing nORFs were identified from human post-mortem brain samples from schizophrenia, control and bipolar disorder patient samples. Read alignment and transcript assembly was done using HISAT2-StringTie and STAR-RSEM as described in the materials and methods section. Around 200,472 transcripts were identified using HISAT2-StringTie and 137922 transcripts using STAR-RSEM. Using the workflow described in **Figure 3**, we identified 3862 and 3617 nORFs within sample transcripts generated using HISAT2-StringTie and STAR-RSEM, respectively. A few example cases of identified nORFs generated using IGV ^43^ are shown in **Figure 5B**. Furthermore, the differences in nORF identification based on the alignment-assembly tool used is presented in **Figure 6**.

**Figure 5:**
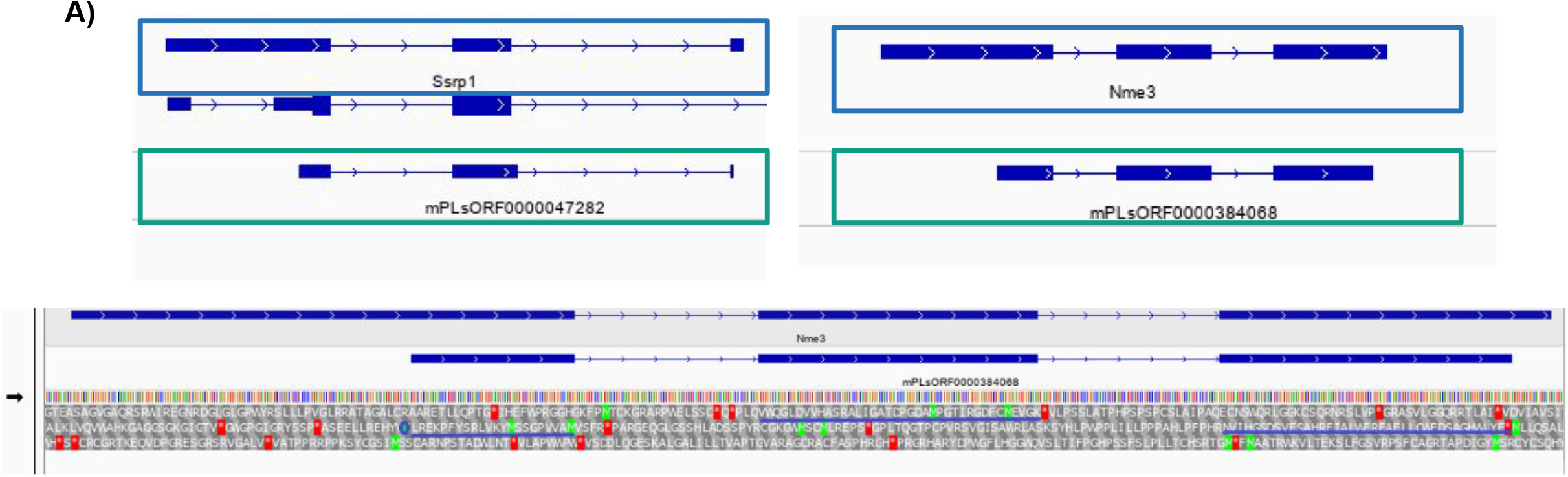
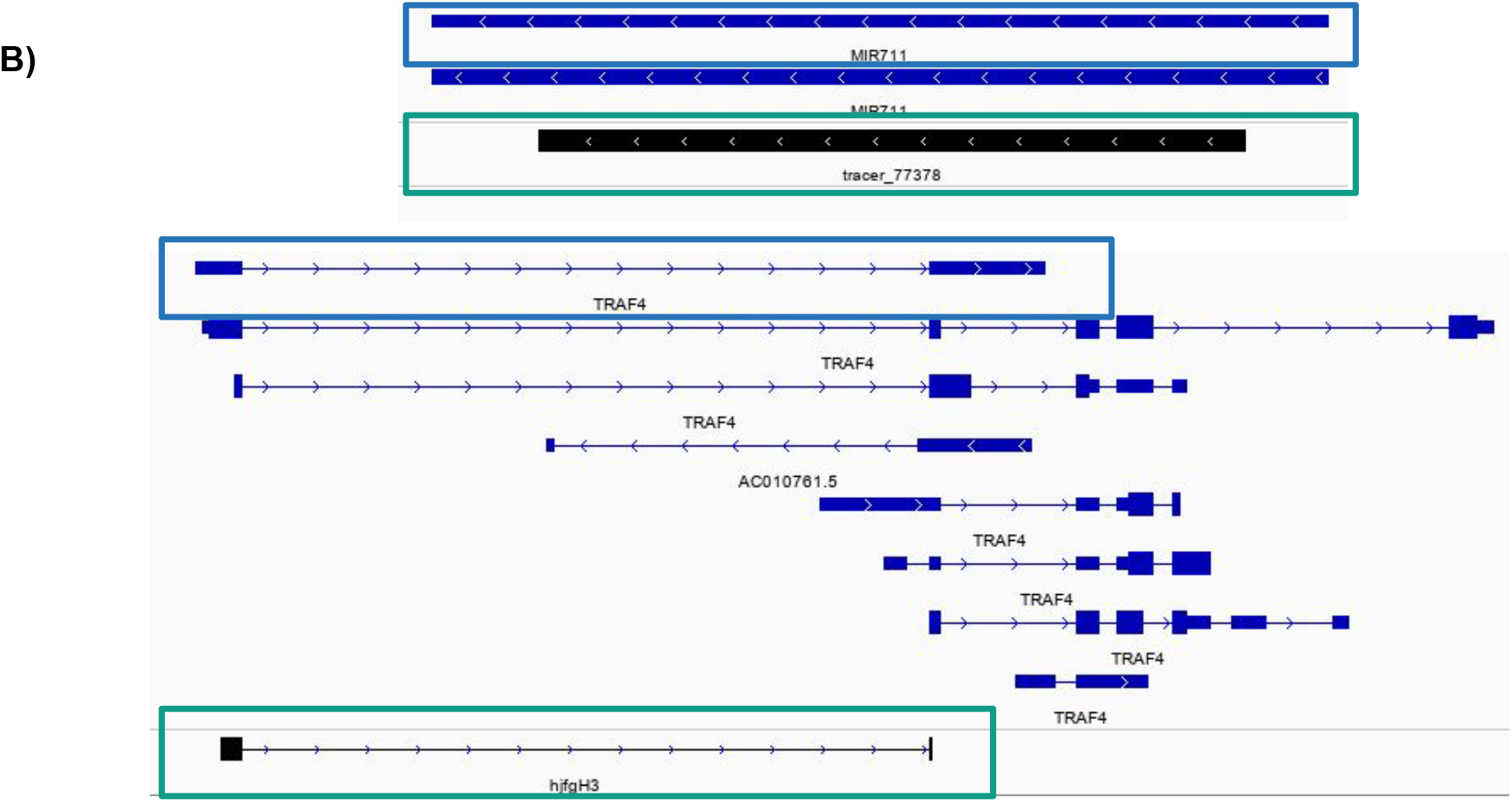
Identification of nORFs at the transcript level visualised using IGV. **(A)** In mouse B and T cells: A nORF (enclosed within the green box) identified within a transcript (enclosed within the blue box) annotated as processed_transcript (top-left) and another within a transcript annotated retained_intron (top-right) are shown. Translation of the nORF (blue lines underlining sequence) within the retained_intron as verified by proteogenomic approach (bottom). **(B)** In neuropsychiatric samples: nORF (enclosed within the green box) identified within a transcript (enclosed within the blue box) annotated as miRNA (top) and another within a transcript annotated retained_intron (bottom) are shown.

**Figure 6:**
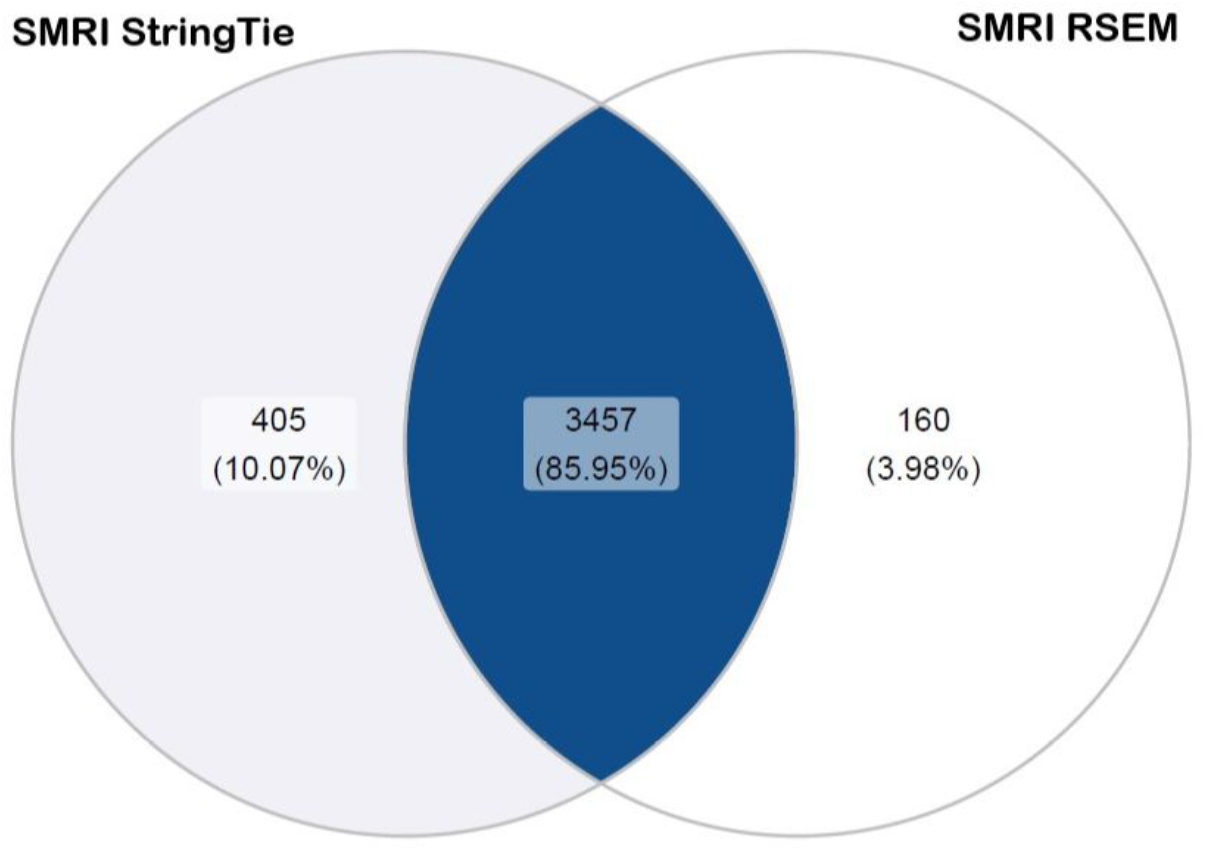
Differences in nORF identification based on read aligner-transcript assembler used for analysis. Of the 200472 and 137922 transcripts assembled by StringTie and RSEM respectively, 3617 and 3526 transcripts were identified with 3862 and 3617 nORFs respectively using the proposed workflow. Approximately, 86% of the total nORFs identified from SMRI samples were called by both the tools whereas StringTie identified an additional 405 nORFs uniquely and RSEM, 160 nORFs uniquely.

## Discussion

The importance and functions of nORFs is still being determined. Usual approaches for nORF identification involve Ribo-Seq data analysis for identifying novel translation events from RNAs or a proteogenomic approach for validating the presence of nORFs within the sample of interest. Although proteogenomics allows for the quantification and study of nORF products, the absence of MS data for most studies, especially those that involve national and international consortiums, such as the TCGA and PsychENCODE, hinders the systematic investigation of this unknown, ever growing repository of protein products with potential functional implications.

In this study we propose a framework for identification of nORF transcripts using available transcript data based on RNA-Seq (and also using Ribo-Seq) for these large-scale datasets. The expression values of nORFs predicted by the framework could then be used for downstream analysis, for example, differential expression analysis between two conditions or correlation of transcript expression with other genic regions of interest. The parameters imposed in the workflow are a bit stringent as we concentrate on nORFs in intronic and intergenic regions, and it is possible that we do not detect several nORFs, for example, nORFs that are embedded within CDS. Our goal is also to minimize ambiguity arising from multiple ORFs and transcripts in the same genomic region and to identify transcripts containing nORFs within them with some certainty.

We show the utility of this workflow by employing it to mouse B and T cells and human neuropsychiatric datasets and demonstrate how nORFs are present within several transcripts conventionally annotated as noncoding. The nORF examples shown in **Figure 5A**, that were identified by the workflow also show evidence of translation^8^. Additionally, we show how different read alignment and transcript assembly tools could lead to the identification of different nORFs because of underlying differences in the assembled transcriptome.

The proposed approach can be used for screening transcriptome of several large-scale studies, without corresponding MS data, for potential novel open reading frames, generating a shortlist of proteins to biochemically validate as described in Prabakaran et al. ^4^. We believe our framework will help in revealing more nORF products from the ‘hidden proteome’ that may shed light on biochemical processes that may not yet have been discovered and possibly disrupted in diseases. This will in turn further call into question the current description of noncoding transcripts and the need to redefine existing genome annotations.

## Acknowledgements

We would like to thank Dr. M. Webster, Dr. Jakub Tomasik and Prof. Sabine Bahn for giving us access to the SMRI dataset, Prof. Erik Miska for giving us access to cichlid RNA-Seq samples and Dr. Matthew Wayland and Adam Boxall for useful discussions during this project. We would also like to thank Cancer Genomics cloud (CGC) for allowing us to use their platform to carry out read alignment and transcript assembly.

## Author contributions

CE did transcriptomic analysis for the B and T cell and brain post-mortem study, contributed to proteogenomic analysis, devised and implemented the proposed workflow and wrote the manuscript. Sh. P did the cichlids analysis, read the manuscript and participated in discussions about the workflow. SP designed and supervised the project, devised the workflow and wrote the manuscript

## Materials and Methods

### Mouse B and T cell samples

Spleen samples from six male and six female C57BL/6J mice were FACS sorted to isolate resting B and naive CD4+ T cells. Resulting total RNA-seq data was processed here. Details of the sample processing can be obtained through Erady et al. ^8^

### Neuropsychiatric samples

23 schizophrenia (SCZ), 16 bipolar disorder (BD) and 23 control (CNT) samples isolated from Brodmann area 46 (BA46) were obtained from the Array collection of SMRI ^44^. Briefly, 1ug of total RNA was poly-A selected using oligo-dT Dynabeads, libraries were prepared using Illumina’s TruSeq v1 (Illumina, Hayward, CA) and sequencing was performed using Illumina HiSeq 2000 giving ~3 Mb of 90bp paired-end reads for each library. Sample reads were assessed for quality using FastQC ^45^ prior to read alignment.

### Cichlid samples

Total RNA-Seq reads depleted with rRNA, from testes and liver tissues of two cichlid fish species *O. niloticus* and *P. nyererei* (n = 3-4) were obtained. Details for sample preparation, RNA-Sequencing and downstream processing steps can be found in Puntambekar et al. ^46^. Parameters for read aligners and transcript assemblers have been described in brief in the following sections.

### Seven Bridges Cancer Genomics Cloud

Read aligners and transcript assembly tools were run on the CGC server (www.cancergenomicscloud.org) ^47^.

### Read aligners and their parameters

#### HISAT2

Neuropsychiatric RNA-Seq reads were aligned using HISAT2 v2.1.0 ^25^, with default parameters except ‘--add-chrname’, ‘—dta’ and ‘--summary-file’ were set to TRUE. Additionally, either Phred +33 or Phred +64 encoding was set to TRUE based on the sample being analysed. Reads were aligned using the index for the GRCh38 genome available at https://ccb.jhu.edu/software/hisat2/manual.shtml.

Cichlid RNA-Seq reads were aligned using HISAT2 v2.1.0 with default parameters. HISAT2 build was used to create the indexed reference using reference genome for the respective fishes downloaded from NCBI (details in Puntamberkar et al. ^46^).

#### TopHat

Cichlid RNA-Seq reads were aligned using TopHat v2.1.0 ^32^ with default parameters using reference genome for the respective fishes downloaded from NCBI.

#### STAR

For neuropsychiatric samples, STAR Genome generate was used to create the required genome files for STAR ^33^ using annotation GTF file and GRCh38 primary assembly FA file from gencode v30 ^48^. STAR v2.6.0c was run on the raw sample read files and the genome files as input with default parameters except −quantMode was set to TranscriptomeSAM.

### Transcript assemblers and their parameters

#### StringTie

For the neuropsychiatric samples, transcript assembly using StringTie v1.3.3 ^25^ was done in three steps for HISAT2 aligned reads. First, StringTie was run with default parameters and ‘-A’ set to TRUE to assemble sample-specific transcripts from the aligned reads (BAM files), using gencode V30 primary comprehensive gene annotation ^48^ as reference. Second, all the GTF files generated in the previous step were merged using StringTie −merge to create a union transcript dataset. Third, StringTie was rerun on the aligned reads with StringTie merged file as the reference and parameters ‘-B’, ‘-e’ and ‘-A’ set to TRUE, allowing us to calculate sample-specific transcript abundances for the union transcript dataset ^25^.

Similarly, for the cichlid samples, StringTie v1.3.3 was run in two different modes: with and without a reference genome, downloaded from NCBI for the respective fishes.

#### Cufflinks

For the cichlid samples, Cufflinks v2.2.1 ^36^ was run with default parameters in two different modes: with reference annotation-based transcriptome assembly and without a reference annotation.

#### Trinity

For the cichlid samples, the *de novo* assembler Trinity v2.0.6 ^37^ was run with default parameters. Additionally, the resulting transcriptome was mapped to their respective genome using GMAP v2017-11-15 ^49^ to extract genomic coordinates of the transcripts for comparison to reference annotation.

#### MAPs

For the neuropsychiatric samples, MAPs ^19^ was run on the BAM files with default parameters and without a reference genome. To generate a union dataset for comparison across samples during differential expression analysis, we fed the results of MAPs to StringTie −merge to create a union transcript dataset. And then, sample-specific transcript abundances were recalculated using StringTie, run with parameters ‘-B’, ‘-e’ and ‘-A’ set to TRUE.

#### RSEM

For the neuropsychiatric samples, RSEM v1.3.1 ^35^ was used for transcript quantification of STAR aligned reads. First, RSEM prepare reference was run on annotation GTF file and GRCh38 primary assembly FA file from gencode v30 ^48^, to generate the required index files. Next, RSEM calculate expression was run with aligner parameter set to STAR, append names, output genome BAM and sort BAM by coordinates set to TRUE.

#### RNA-Seq read simulation for cichlids

To decide upon a transcript assembler and assess the precision and sensitivity of *de novo* and reference-based transcriptome assembly, RNA-Seq reads from *O. niloticus* and *P. nyererei* were simulated using the R package polyester v1.14.1 ^50^. Three replicates of ~25 million 75bp paired-end reads were simulated for each species without sequencing errors and with uniform transcript expression levels. Simulated reads were aligned and assembled into transcripts as described in the sections above.

#### Precision and sensitivity analysis of simulated cichlid transcriptomes

The precision and sensitivity of the simulated transcriptomes obtained from multiple transcript assemblers, were determined with respect to the reference annotation. In brief, 10 x 10,000 transcripts were randomly sampled with replacement from each simulated transcriptome and compared to the reference transcriptome using GffCompare v0.10.1 ^42^. The precision and sensitivity estimates obtained from GffCompare were then used for further analysis. Additionally, since we use a subset of the data here which can lead to loss of sensitivity, raw sensitivity values were multiplied by transcriptome size/1000 to account for this loss.

## Supplementary Figures

**Supplementary Figure 1:**
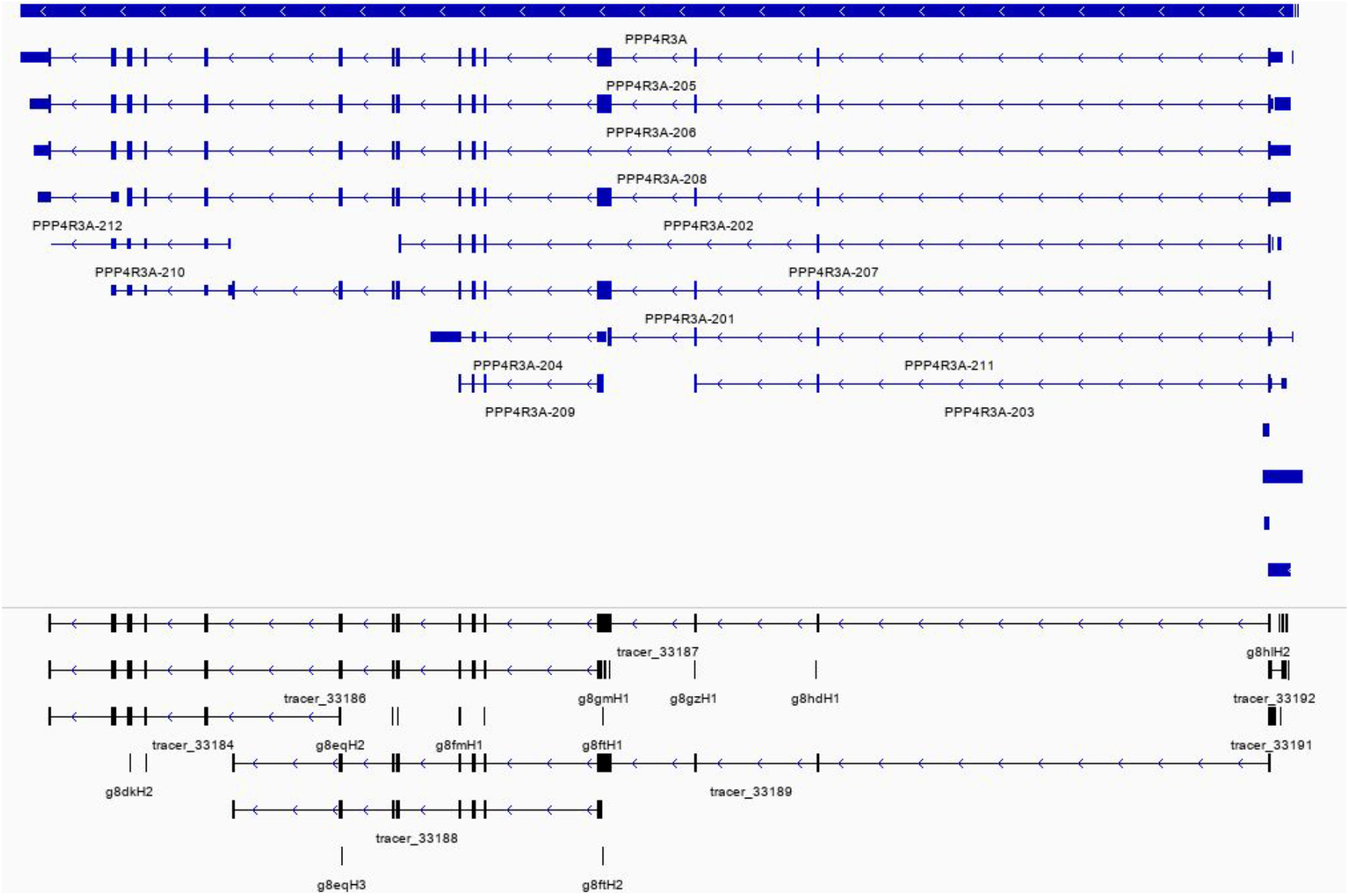
Challenges in nORF-transcript identification. An example of protein phosphatase 4 regulatory subunit 3A (PPP4R3A) gene along with its transcripts (in blue) along with the different nORFs mapping to this region (in black) visualised using IGV.

**Supplementary Figure 2:**
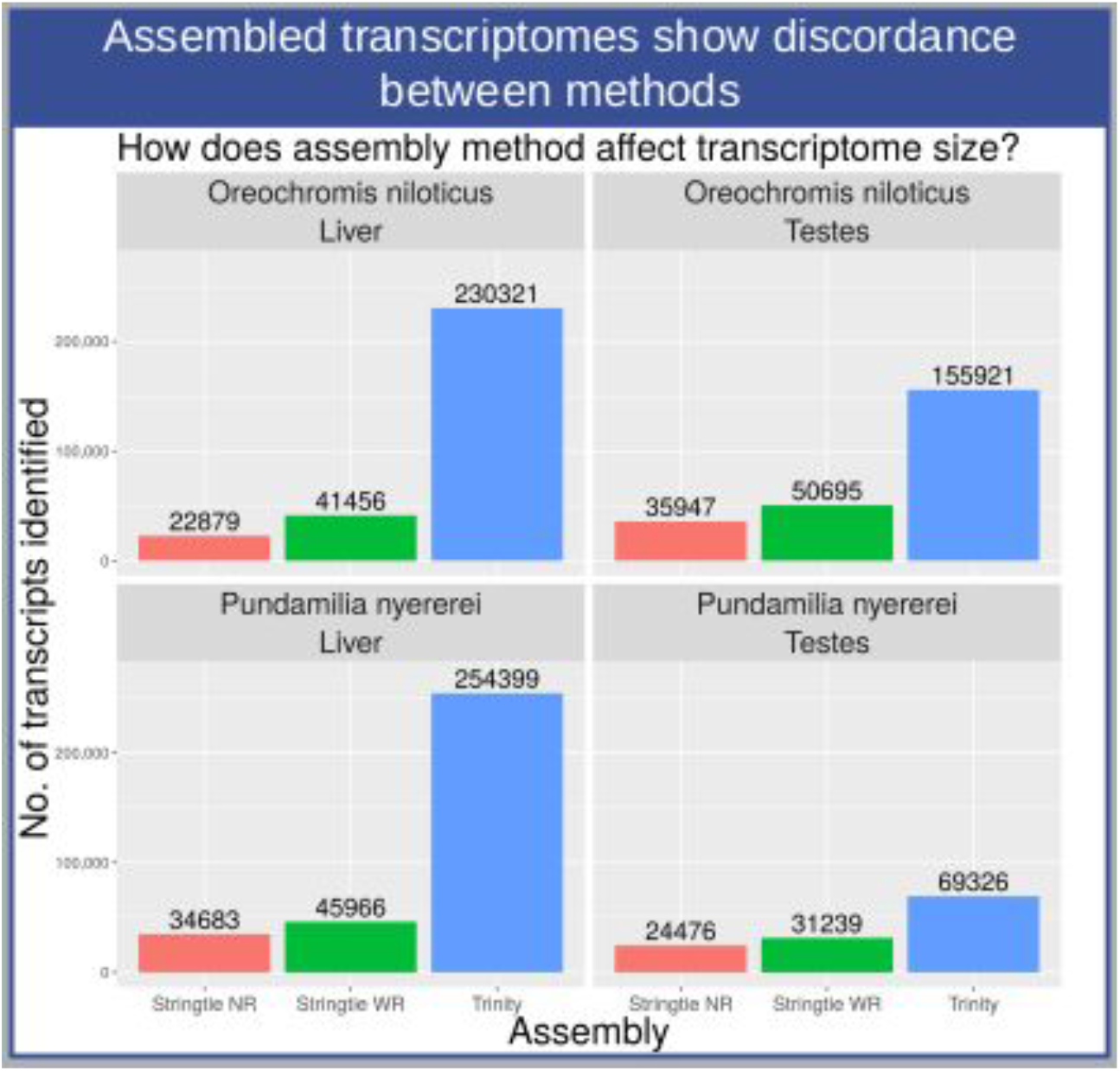
Comparison of different transcript assembly tools. Transcripts were generated using StringTie with reference genome (StringTie WR), StringTie without reference (StringTie NR) and Trinity. Depending on the tool used different transcript sets are identified with Trinity identifying a much larger proportion than StringTie run in de novo mode. But since the sensitivity and precision of Trinity generated transcripts is much lower than that of StringTie (Supplementary figure 2), we decided to not work with it.

**Supplementary Figure 3:**
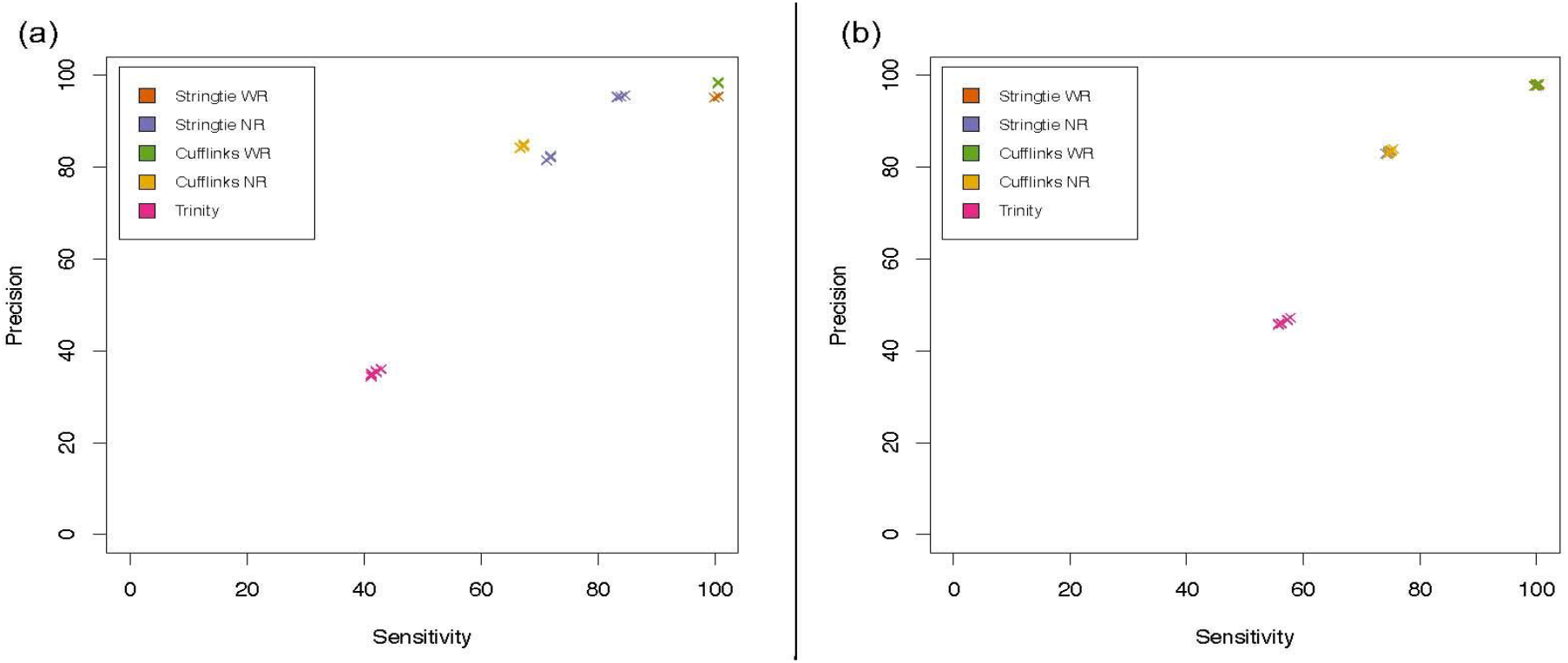
Sensitivity and precision of transcriptome assembly of simulated reads. Simulated reads with uniform expression levels and no sequencing errors were assembled using five transcriptome assembly methods. 10×10,000 transcripts were randomly sampled with replacement from each simulated transcriptome and the sensitivity and precision of these subsets assessed using GffCompare. (a) O. niloticus-derived reads. (b) P. nyererei-derived reads. Trinity had the lowest sensitivity and precision scores.

